# Phylogenetically-informed crayfish conservation in the face of climate change

**DOI:** 10.1101/2025.06.03.657651

**Authors:** Sebastian Pipins, Monika Böhm, Lucie Bland, Keith A. Crandall, Md Anwar Hossain, James Rosindell, Rikki Gumbs

**Author notes:** Corresponding author. E-mail address (S. Pipins).

## Abstract

Crayfish are an ancient clade of freshwater decapods that play vital and diverse ecological roles in the freshwater systems that they inhabit. One third of assessed crayfish species are threatened with extinction and 87% are highly sensitive to climate change. However, the extent to which the evolutionary history of crayfish is threatened, especially by climate change, remains unclear. To address this, we produced a phylogenetically-informed species prioritisation for the conservation of crayfish and explored the consequences of projected climate change scenarios on crayfish phylogenetic diversity. We used the EDGE2 metric to highlight 70 priority species for conservation. Their combined threatened evolutionary history accounts for 40.7% of the expected phylogenetic diversity loss for all 673 crayfish. Of these priority species, 45.7% are vulnerable to climate change, with the majority (32 species) located in Australia. Under a high warming scenario, individual species that are more evolutionarily distinct are projected to be disproportionately affected. The extinction of climate change vulnerable species could cause the loss of up to 26.3% more phylogenetic diversity than the random expectation, equivalent to 409 million years of evolutionary history. Our results highlight the need to consider climate change effects on phylogenetic diversity to fully understand its impact on biodiversity. If actions were taken to avoid a worst-case warming scenario in favour of an intermediate scenario, we could safeguard 251 million years of unique crayfish evolutionary history.

## Introduction

Despite covering less than 1% of the Earth’s surface, freshwater ecosystems contain approximately one tenth of all recorded animal species (Balian et al., 2007). Freshwater biodiversity is vital for human health and well-being, but these ecosystems are in a state of crisis (Albert et al., 2020) due to a myriad of anthropogenic stressors (Dudgeon et al., 2006; Vörösmarty et al., 2010). Superimposed on these is the growing threat of climate change (Reid et al., 2019), which could threaten half of all freshwater fish globally (Darwall & Freyhof, 2016). Monitored populations of freshwater vertebrate species have declined on average by 85% since 1970 (WWF, 2024), a decline notably higher than their terrestrial or marine counterparts.

Crayfish comprise more than 690 species worldwide (Crandall & De Grave, 2017; DecaNet, 2024), split across three northern hemisphere families (Astacidae, Cambaridae, Cambaroididae) and one southern hemisphere family (Parastacidae). They play several important roles in freshwater communities (Reynolds et al., 2013), including as keystone species (Füreder et al., 2010), ecosystem engineers (Harvey et al., 2011), indicators of water quality (Pârvulescu et al., 2011) and some, like the noble crayfish (Astacus astacus), serve as umbrella species for the conservation of others (Edsman et al., 2010; Reynolds et al., 2013). Crayfish have been the subject of IUCN Red List assessments (Richman et al., 2015) and, together with other freshwater decapods, fish, and odonates, they are part of a global effort to assess the status of the world’s freshwater fauna (Sayer, 2024). Around one-third of assessed crayfish are threatened with extinction, with European crayfish experiencing the greatest number of different threats and Australian species being most at risk from climate change (Richman et al., 2015).

To conserve crayfish biodiversity into the future, certain species and regions will have to be prioritised. One way of directing conservation efforts is to consider phylogenetic diversity (PD; Faith 1992), which is calculated as the sum of all branch lengths connecting a set of species on a phylogenetic tree (Faith, 1992). The conservation of species without consideration of their evolutionary heritage can lead to substantial losses from the tree of life (Gumbs et al., 2023a). Therefore, prioritising PD not only helps to capture variety among living organisms (Faith, 2008), but also deals with the issues of limited resources by highlighting priority species or areas for conservation that conserve evolutionary history (Crandall, 1998; Whiting et al., 2000). Recent work has also empirically demonstrated that maximising PD captures a greater proportion of diverse features that are beneficial to humanity (such as food and medicine) compared to the random selection of taxa (Molina-Venegas et al., 2021).

The PD conservation framework prioritises species that both represent a disproportionately large amount of evolutionary history and are threatened with extinction (Isaac et al., 2007; Mooers et al., 2008). One way of doing so is through the EDGE metric, which ranks species based on a combination of their irreplaceability (ED: Evolutionary Distinctiveness) and their vulnerability (GE: Global Endangerment) (Isaac et al., 2007). The metric has been applied to several taxonomic groups spanning vertebrates (Gumbs et al., 2018; Isaac et al., 2012; Isaac et al., 2007; Jetz et al., 2014; Stein et al., 2018), invertebrates (Curnick et al., 2015), and plants (Forest et al., 2018). More recently, the method for prioritising species has been updated (Gumbs et al., 2023b); the ‘EDGE2’ metric is now better able to incorporate uncertainty in phylogenetic and extinction risk data and partially weights a species’ value by the extinction risk of related species (Faith, 2008; Steel et al., 2007). The EDGE2 metric has since been applied to all jawed vertebrates (Gumbs et al., 2024), but so far no invertebrate group has been assessed despite the metric’s greater capacity to deal with data-poor clades.

Recent research has led to a number of exciting developments for crayfish, including descriptions of new species, taxonomic revisions (Crandall & De Grave, 2017), the production of a comprehensive molecular phylogeny (Stern et al., 2017), and a trait-based climate change vulnerability analysis (Hossain et al., 2018). Whilst we know that climate change is predicted to confer significant changes in the PD of some clades such as eucalypts (Gonzalez-Orozco et al., 2016), we do not know how it will affect the evolutionary history of most other groups, such as crayfish. In this study, we therefore determined the extent of crayfish PD threatened by climate change and the PD that could be saved by avoiding a high warming scenario. We also produced a priority EDGE list for the conservation of crayfish. Our findings highlight the most irreplaceable and vulnerable crayfish globally, with the potential to guide future conservation action in the face of climate change.

## Materials and Methods

### EDGE2

EDGE2 scores were calculated by assigning each species a probability of extinction (a GE2 score) from a distribution of possible values corresponding to its IUCN Red List category (Gumbs et al., 2023b). The median values of each distribution are as follows: Least Concern = 0.060625, Near Threatened = 0.12125, Vulnerable = 0.2425, Endangered = 0.485, Critically Endangered = 0.97. The extinction risk probabilities for Data Deficient or unassessed species are drawn from a pool of all possible values, with the median value being roughly equivalent to the Vulnerable Red List category. We believe this is an appropriate representation given the link between data-poor species and elevated extinction risks (Bland et al., 2015; Borgelt et al., 2022; Howard & Bickford, 2014).

The branch lengths of a phylogenetic tree are then multiplied by the product of the probabilities of extinction for all descendant species to estimate the expected PD loss for each branch (Gumbs et al., 2023b). Summing the branch lengths that connect a species from the tip to the root of the phylogeny gives its EDGE2 score, which is a measure of the amount of threatened evolutionary history a species is responsible for. ED2 scores are calculated in the same way but preclude the focal species’ probability of extinction from the calculation. Summing all branches of an extinction risk-weighted tree gives the total expected PD loss for a given clade. The EDGE2 calculation accounts for phylogenetic complementarity, meaning that the extinction probabilities of related species will influence each other’s scores (Steel et al., 2007). EDGE2 scores therefore give greater precedence to species on long, isolated branches of a phylogenetic tree, or to threatened and unique species whose closest relatives are also threatened.

To calculate EDGE2, we first revised the most comprehensive crayfish phylogeny to date (Stern et al., 2017) by inserting known missing species following the methods described in Gumbs, et al. (2023b). We identified missing species based on the crayfish revision of Crandall & De Grave (2017). There were 144 differences between the tip labels in Stern et al.’s phylogeny and the species listed in Crandall & De Grave, largely a result of recent taxonomic revisions to certain groups and new species descriptions. After matching the taxonomies and removing species not listed in Crandall & De Grave, the Stern et al. phylogeny was left with 435 species tips. We imputed missing species into the phylogenetic tree using the function congeneric.impute in the pez package in R (version 3.6.1) (Pearse et al., 2015), which inserts missing species to a random site within the crown group of the lowest taxonomic rank available (genus, family etc.). At least 23 new species of crayfish have been described since Crandall & De Grave (2017), including 17 species in the family Cambaridae (Bloom et al., 2019; Foltz et al., 2018; Glon et al., 2020; Glon et al., 2019a; Glon et al., 2019b; Johnson, 2018; Loughman et al., 2019; Loughman & Williams, 2018; Lyko, 2017; Pedraza-Lara et al., 2021; Perkins et al., 2019; Taylor, 2018; Álvarez et al., 2021), five in Parastacidae (Huber et al., 2018; Huber et al., 2020; Lukhaup et al., 2018; Patoka et al., 2017), and one in Astacidae (Pârvulescu, 2019). These species were therefore also imputed into the phylogeny. To capture the uncertainty in the placements of imputed species, we conducted the imputation 1000 times.

We then calculated EDGE2 scores for all 673 crayfish species using Red List data (Version 2019-3) (IUCN, 2019), which returned categories for 590 species. We calculated PD and expected PD loss for crayfish to determine the extent of threatened crayfish evolutionary history. The median terminal branch lengths (TBL) of threatened and non-threatened crayfish were contrasted using a Mann Whitney U test. For all Mann Whitney U tests, we used Cliff’s delta d to quantify effect size (Cliff, 1993). We then determined the phylogenetic signal of extinction risk in crayfish using Pagel’s λ by using species probability of extinction scores across the 1000 possible trees and taking the median value. Using Analysis of Variance (ANOVA) with Tukey’s Honest Significant Difference tests, we then compared the TBL and EDGE2 scores of crayfish with tetrapod animal clades using data from Gumbs et al. (2024).

### EDGE species

Species are identified as priority EDGE species if they are in a threatened IUCN Red List category (Vulnerable, Endangered, Critically Endangered) and have a median EDGE2 score that is above the median for all species in at least 95% of iterations (Gumbs et al., 2023b). We used IUCN Red List data to map EDGE crayfish by country. We further determined the extent of expected PD loss contributed solely by EDGE crayfish by pruning the extinction risk-weighted phylogenies to contain only EDGE species and summing the branch lengths. Similarly, we also estimated the proportion of expected PD loss contributed by EDGE species. To compare if the contribution of EDGE species to total expected PD loss is consistent across clades, we repeated this procedure using mammal EDGE data (Gumbs et al., 2023b).

In line with the EDGE2 framework, we categorized species into three additional groups: an EDGE2 Watch List, for species that have EDGE2 scores above the median in 95% of cases but are listed as either Least Concern or Near Threatened; an EDGE2 Research List, for species with EDGE2 scores above the median in 95% of cases and that are either Data Deficient or unassessed by the IUCN; and an EDGE2 Borderline List, for threatened species whose EDGE2 scores are above the median for the clade in 80% of iterations.

### Data uncertainty

The EDGE2 methodology built on the original EDGE metric (Isaac et al., 2007) by incorporating uncertainty in both phylogenetic and extinction risk; the original EDGE metric (Isaac et al., 2007) could only compute scores for species with sufficient data. To test how this methodological improvement affects species priority lists, we compared the species ranks obtained via the EDGE2 metric and the original EDGE metric for the 446 species with both phylogenetic and Red List data using a Spearman’s correlation. The EDGE metric combines a species’ evolutionary distinctiveness (ED) with its extinction risk (GE) and is calculated as EDGE = ln(1 +ED) + (GE x ln(2)) (Isaac et al., 2007).

A phylogenetic prioritisation was previously applied to crayfish (Owen et al., 2015) using the software ‘Mesquite’ (Maddison, 2005; Maddison & Mooers, 2007), which differs from the original EDGE metric in that the ED score is multiplied by a probability of extinction rather than being summed with the GE index. We used the probabilities of extinction calculated in this study to run a Spearman’s correlation test between Mesquite EDGE, Isaac et al. EDGE, and EDGE2 to determine the consistency of the methods.

We then explored the effect of species imputation on EDGE2 priority lists by comparing how the scores and ranks of the original set of species from the Stern et al. phylogeny differed with and without the imputation of missing species. To do this, we conducted a series of Spearman’s correlations to compare EDGE2 scores, ED2 scores and species ranks of the original set of species before and after phylogenetic imputation. We also calculated the expected PD loss for both sets of species.

### Climate change

To determine the extent of crayfish PD threatened by climate change, we compared the amount of PD lost between species vulnerable to climate change and a null model of random extinction. Climate change vulnerable (CCV) species were those identified by a recent traitbased vulnerability analysis for crayfish (Hossain et al., 2018) focusing on species traits relating to sensitivity, adaptive capacity, and exposure to climate change (Foden et al., 2013). In our current study, we considered two of the warming projections investigated in Hossain et al. (2018), intermediate and high (relating to RCP6.0 and RCP8.5, respectively), at two points, 2050 and 2070. Across scenarios, a total of 126 different species were listed as vulnerable to climate change; under an intermediate warming scenario in 2050 and 2070 there were 87 and 94 CCV species, respectively, whilst there were 107 and 112 species CCV under a high warming scenario (Hossain et al., 2018). We assessed the phylogenetic signal of climate change vulnerability in crayfish using the D statistic (Fritz & Purvis, 2010).

To simulate all CCV species going extinct, we pruned the CCV species for each scenario from each of the 1000 trees and calculated the mean reduction in PD. We then repeated this procedure using a random sample of species of equal size to that of each set of CCV species and where the sets were drawn afresh for each of the 1000 iterations. We tested for differences in PD loss between observed CCV species and randomly drawn species using Mann Whitney U tests. To determine whether climate change is set to affect more evolutionarily distinct species in general, we compared the ED2 scores of CCV species to random sets of species of an equivalent size using Mann Whitney tests.

Finally, we compared the simulated PD loss and ED2 scores of crayfish listed as vulnerable to climate change by Hossain et al. (2018) to those recorded as threatened by climate change on the Red List using Mann Whitney U tests.

## Results

### Crayfish evolutionary history

Across 673 species, crayfish represent 14,934 million years (MY) of phylogenetic diversity, of which 18.5% (2,769 MY) is at risk of being lost. EDGE2 scores ranged from 0.02 MY to 63 MY (Fig. 1), with the median being 1.51 MY. The median TBL of a crayfish species is 8.46 MY (ranging from 0.157 – 126 MY) and is higher in threatened species (10.2 MY) than non-threatened species (8.11 MY; Mann Whitney U: 41720, p = < 0.05), though the effect size was small (d = 0.118). Extinction risk displayed a weak phylogenetic signal in crayfish (λ = 0.161).

**Fig. 1.**
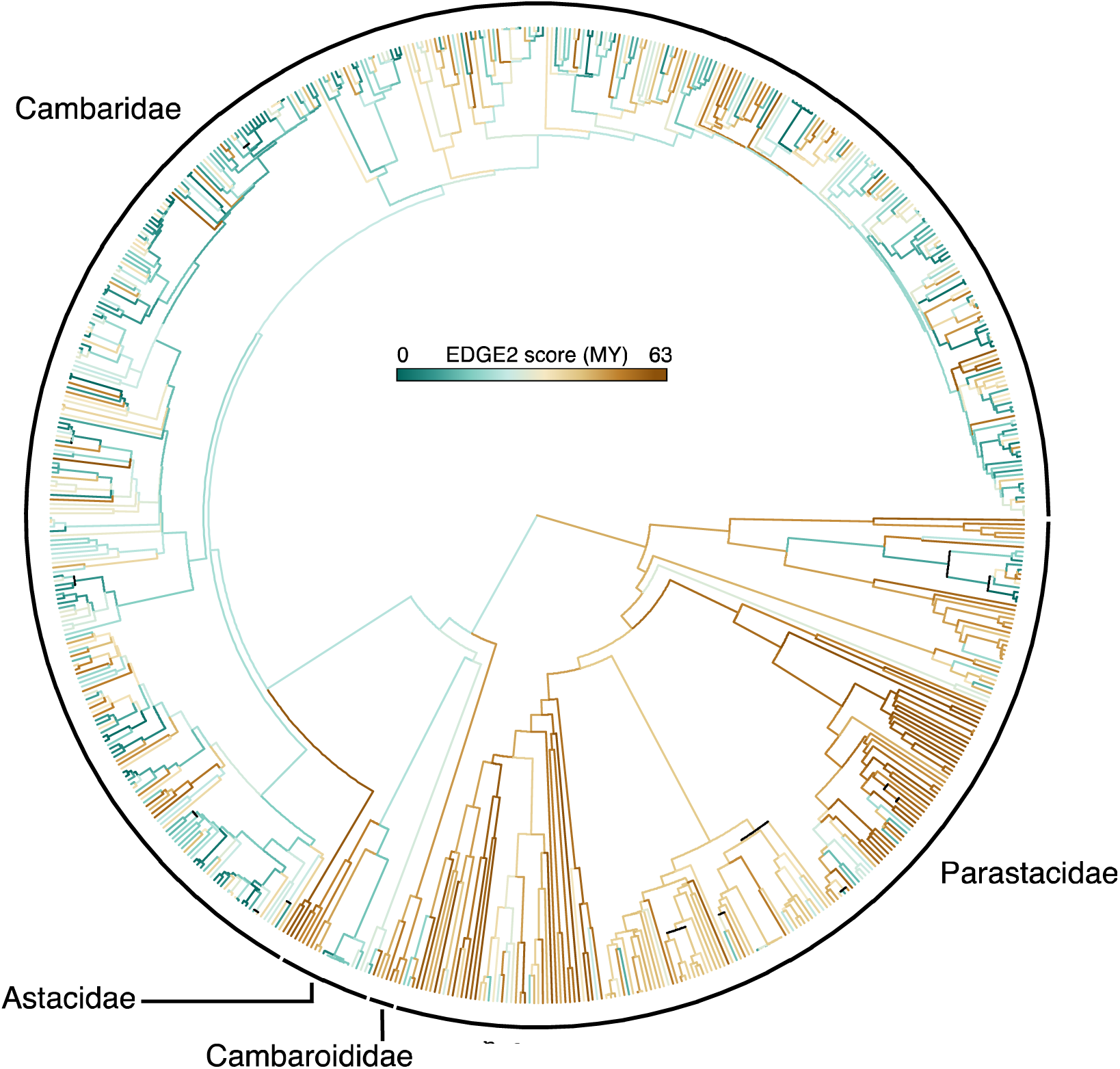
A crayfish phylogeny coloured by EDGE2 score, with higher values in brown and lower values in green, and with crayfish families labelled.

ANOVA with Tukey’s Honest Significant Difference test showed that the median log TBL and EDGE2 score of crayfish is significantly greater than observed in amphibians, squamates, birds, and mammals (p < 0.0001 for all comparisons; Fig. 2). Turtles showed higher log TBL and EDGE2 scores than crayfish (p < 0.0001), whilst crocodilians showed greater log EDGE2 scores (p < 0.01) and non-significantly different log TBL.

**Fig. 2.**
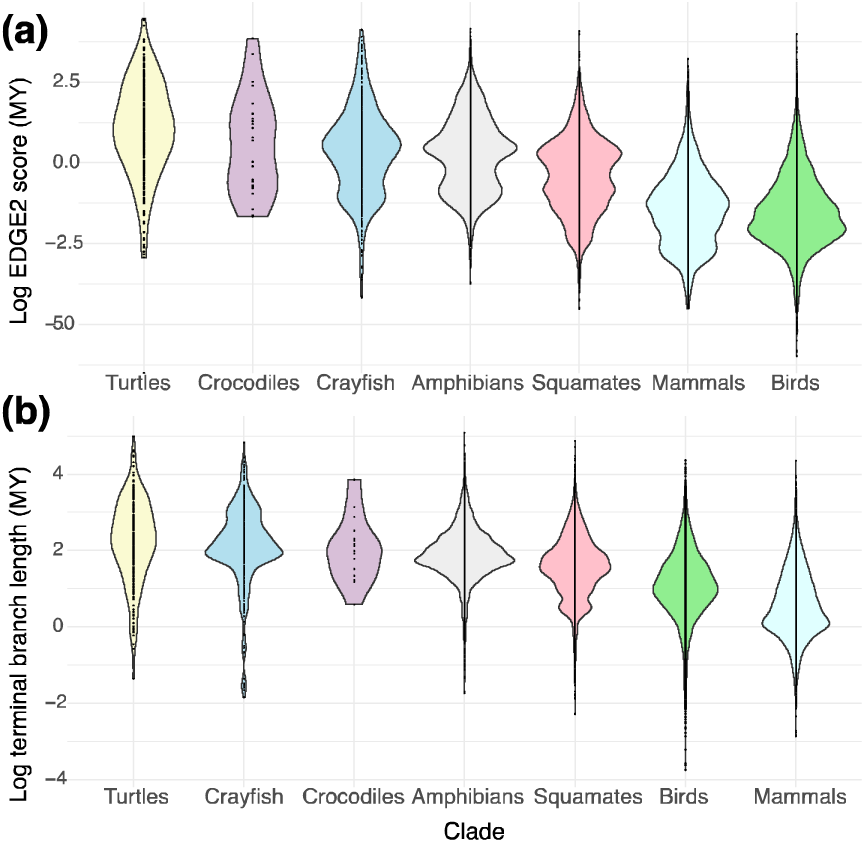
Crayfish log terminal branch length and EDGE2 scores compared with tetrapod animal clades.

There was a significant positive correlation between the original EDGE and EDGE2 ranks for the 446 species with data sufficient IUCN assessments (ρ = 0.929, p < 0.0001, r^2^ =.863), meaning those species ranked as high priority in EDGE were also likely to be ranked highly in EDGE2 (Appendix S1). Furthermore, Mesquite EDGE ranks were strongly correlated with the original EDGE metric ranks (ρ = 0.995, p < 0.0001, r^2^ =.99) and with EDGE2 ranks (ρ = 0.922, p < 0.0001, r^2^ =.85).

### EDGE Crayfish

Seventy species were in a threatened IUCN Red List category and had an EDGE2 score above the median with 95% certainty and were thus deemed EDGE species (Table 1; see Figshare repository).

**TABLE 1.**
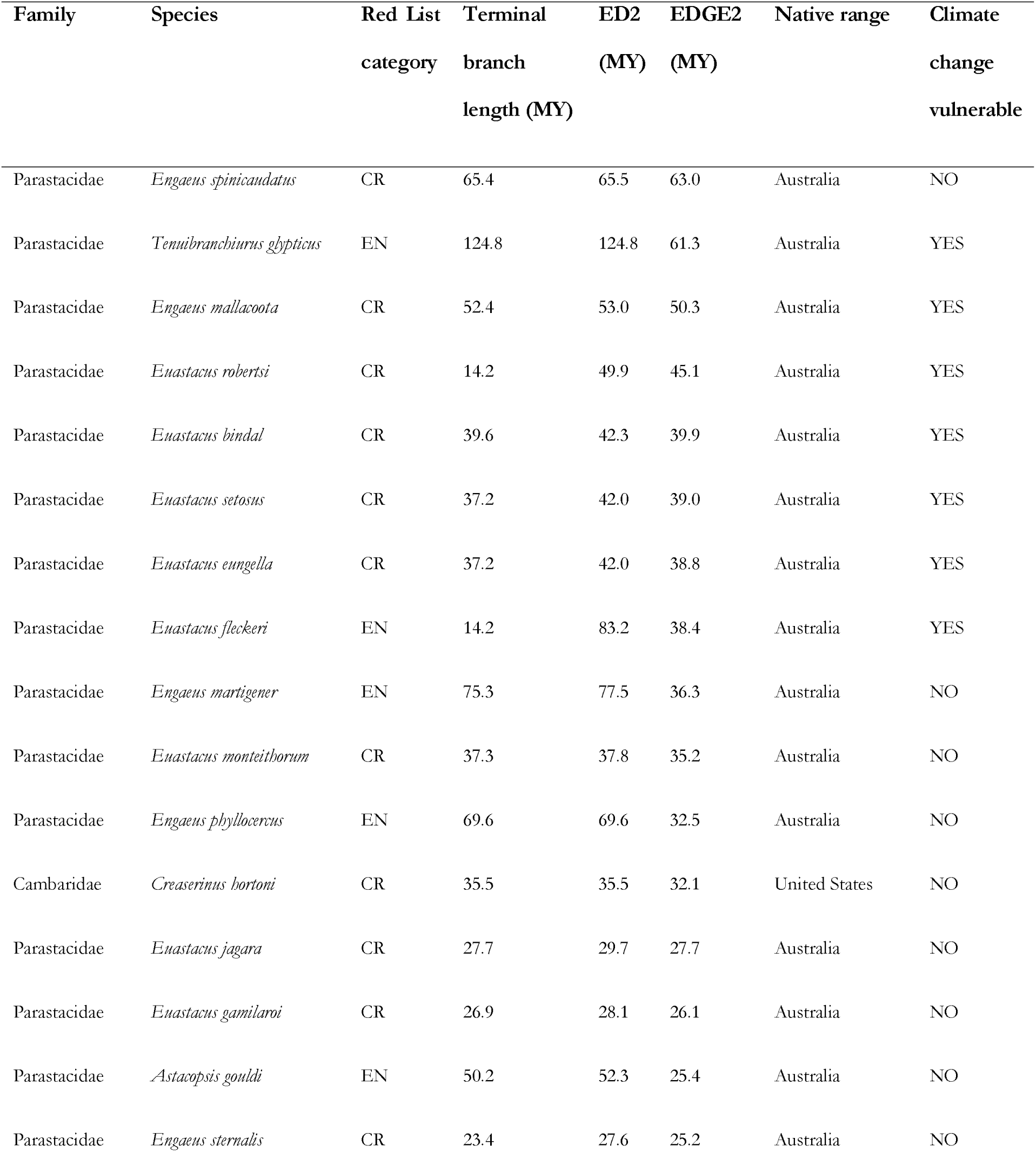
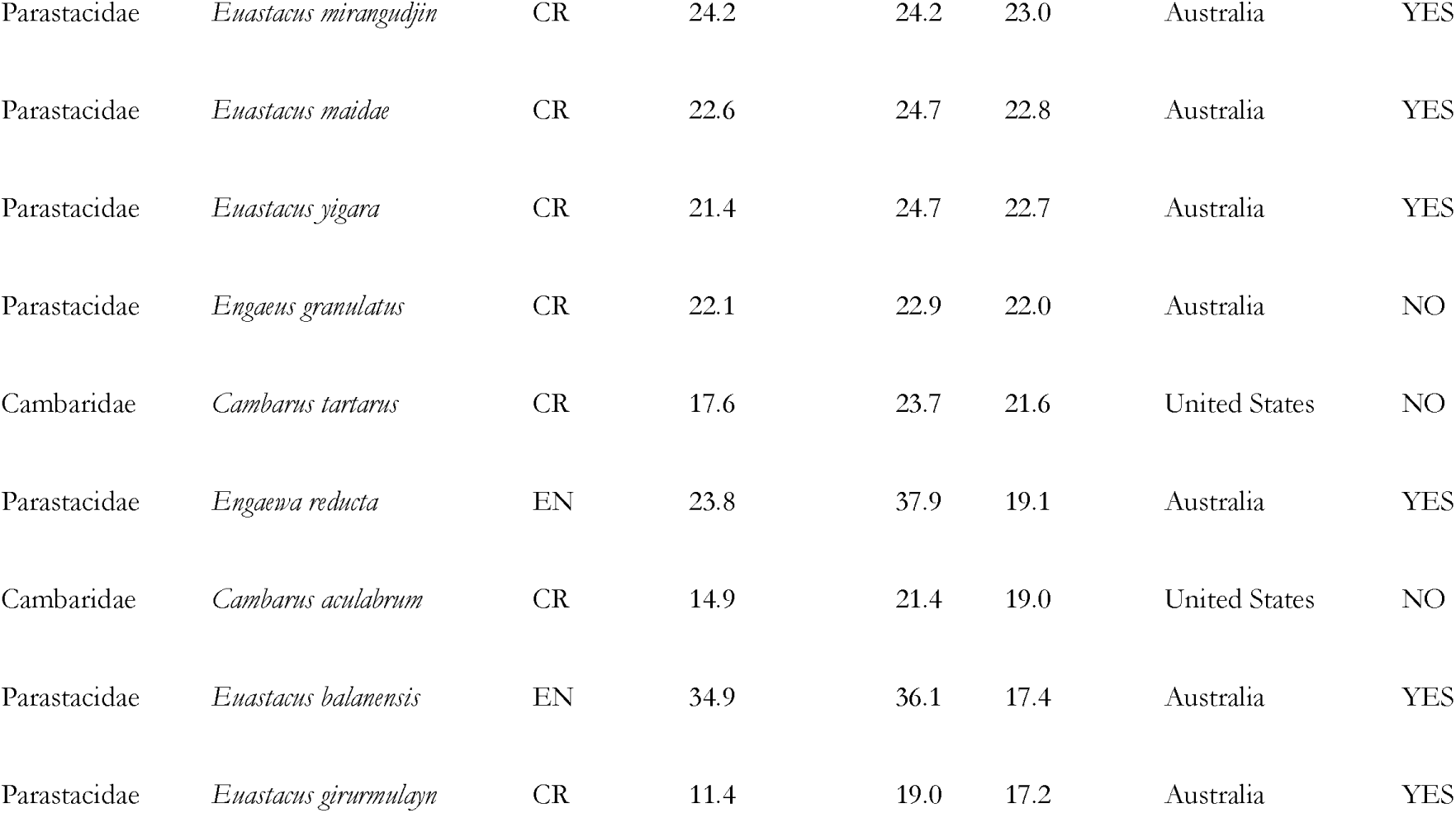
The top 25 EDGE crayfish species, where climate change vulnerability (marked as YES/NO) was taken from Hossain et al. (2018).

These species had EDGE2 scores ranging from 2.75 to 63 MY (median = 13.8 MY), and span 14 genera, three families, and four main regions, with 44 species in Australia, 21 in USA, four in Mexico, and one, Austropotamobius pallipes (the white clawed crayfish), found in Europe (Fig. 3). There were 29 Critically Endangered species, 34 Endangered species, and 7 Vulnerable species. The top ranked EDGE species was the Critically Endangered Scottsdale burrowing crayfish, Engaeus spinicaudatus, a Tasmanian endemic.

**Fig. 3.**
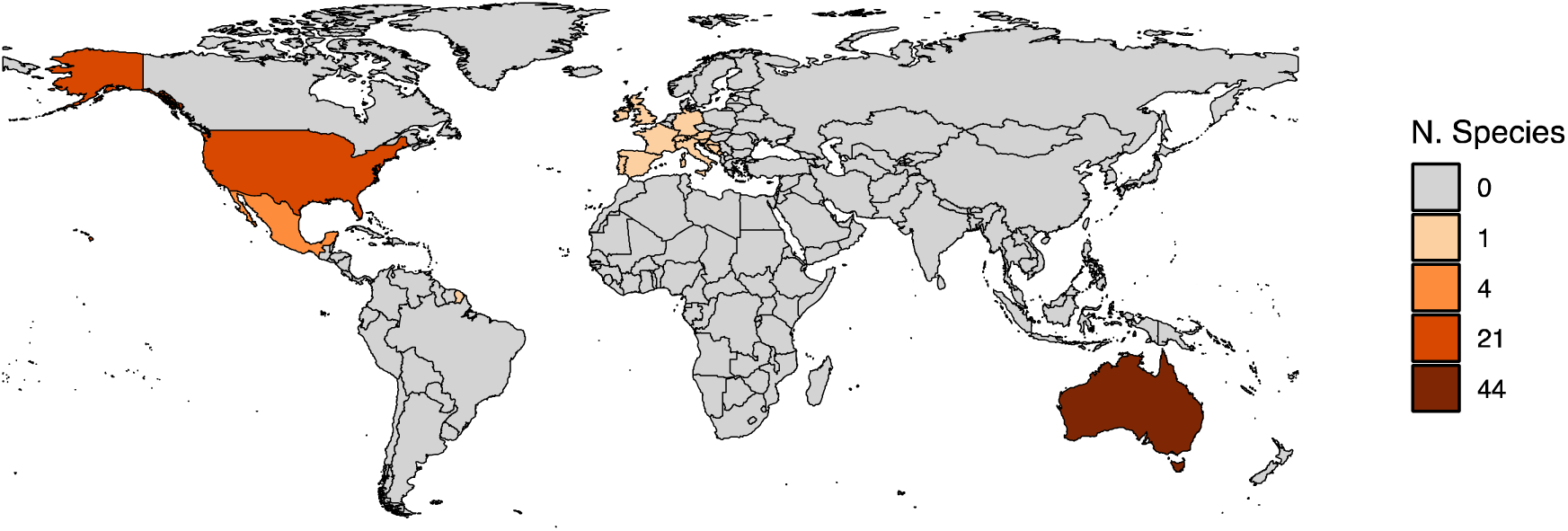
The number of priority EDGE crayfish per country, where presence in a country was determined using IUCN data.

Of the 2,769 MY of crayfish PD that is expected to be lost given present day extinction risks, 40.8% (1,128 MY) is contributed by the 70 EDGE crayfish, despite EDGE species representing just 10.3% of species richness. This is comparable to that observed in mammals, where 36% of the expected loss of the clade is attributable to 10.3% of species. Together, the 140 threatened crayfish species represent 52% (1,440 MY) of expected crayfish phylogenetic diversity loss, of which 78.5% of this is attributable to the 70 that are EDGE listed.

Seven non-threatened species had EDGE2 scores above the median with 95% confidence; one, the monotypic Samastacus spinifrons (endemic to Argentina and Chile), is Data Deficient and was therefore placed on the EDGE Research List. The six others were all Near Threatened and were therefore placed on the EDGE Watch List. Thirty-three threatened species had EDGE2 scores above the median with 80% confidence and were therefore marked as Borderline EDGE species. EDGE2 scores, EDGE lists and expected phylogenetic diversity loss trees will be made available on Figshare on publication (10.6084/m9.figshare.27283635).

### Data uncertainty

Overall, 227 species (33.7%) lacked extinction risk data (i.e., they were Data Deficient or not evaluated on the IUCN Red List) and 238 (35.4%) were without phylogenetic information. Not accounting for these species would lead us to consider 20.7% (3,088 MY) less phylogenetic diversity and 28.6% (791 MY) less expected loss, thus dramatically underestimating the true extent of crayfish threatened evolutionary history.

A Spearman’s correlation revealed that species ranks (Sr = 0.979, p <.0001, r^2^ =.959), EDGE2 scores (Sr = 0.979, p <.0001, r^2^ =.959), and ED2 scores (Sr = 0.973, p <.0001, r^2^ =.902) for the 435 species featuring in the Stern et al.. phylogeny were highly consistent before and after the imputation of missing species (Fig. 4).

**Fig. 4.**
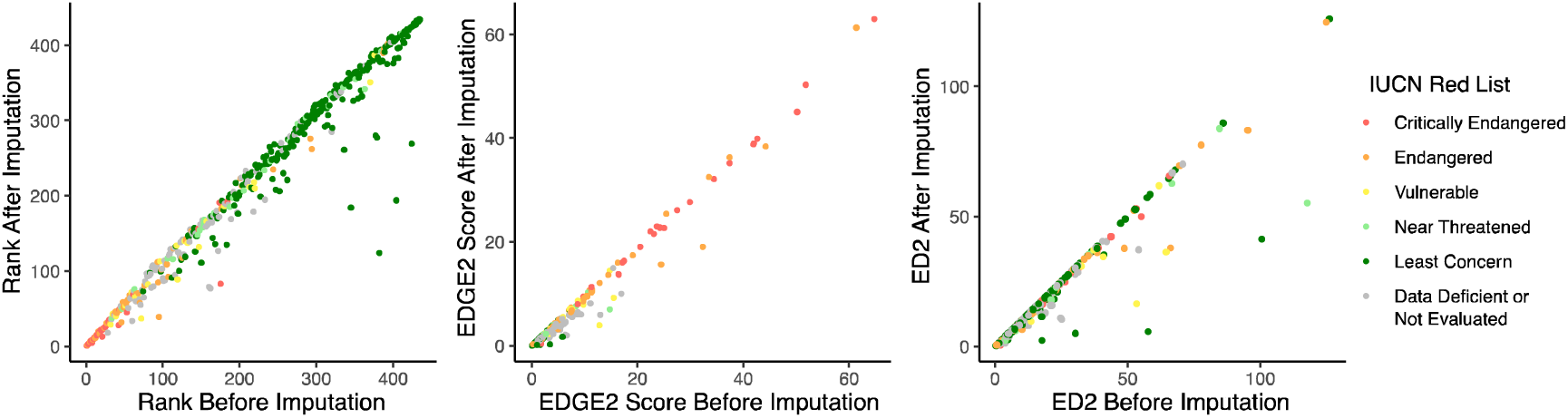
The effect of phylogenetic imputation on species ranks, EDGE2 scores, and ED2 scores. Points represent individual species, coloured by their IUCN Red List categories.

### Climate change

For all four scenarios explored, the simulated extinction of climate change vulnerable species represents a greater loss of total PD than random groups of the same size (Fig. 5). Crayfish projected to be CCV under a 2070 high warming scenario (Hossain et al., 2018) showed the greatest departure from the random expectation, threatening 409 million more years of evolutionary history (Mann Whitney U = 860578, p < 0.0001, d = 0.721), followed by 315 MY of PD loss for crayfish that are CCV under a 2050 high warming scenario (Mann Whitney U = 804058, p < 0.0001, d = 0.608). Both groups of crayfish projected to be CCV under an intermediate warming scenario in 2050 and 2070 showed a similar mean difference in PD loss of 158 MY and 152 MY, respectively (Mann Whitney U = 657521, p < 0.0001, d = 0.315; Mann Whitney U = 652865, p < 0.0001, d = 0.306). The extinction of CCV crayfish species therefore represents between 11.9% and 26.3% more PD being lost than the random expectation.

**Fig. 5.**
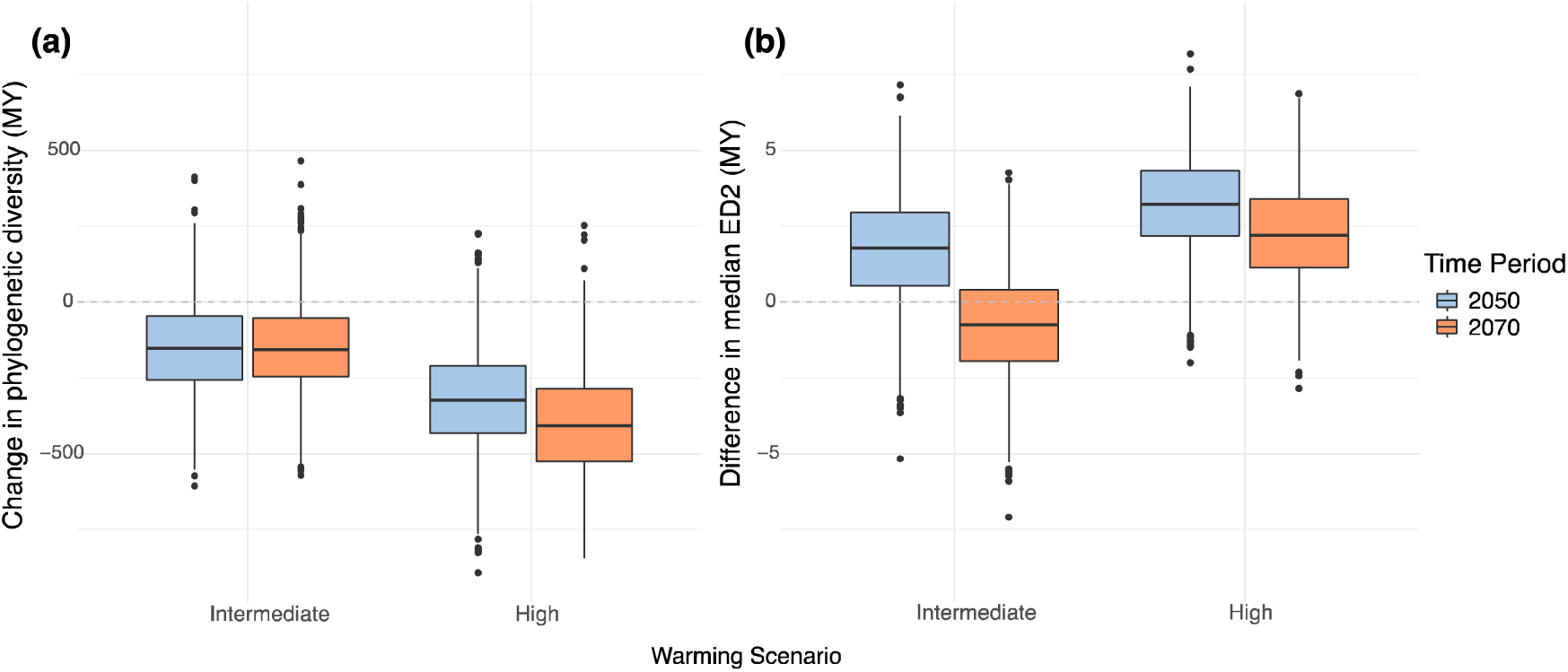
The effect of climate change on crayfish evolutionary history. Left: The median change in phylogenetic diversity resulting from the loss of all climate change vulnerable (CCV) species beyond that of the random expectation (denoted by the black dotted line) under each warming scenario, where negative values represent a greater loss. Right: The median ED2 of a CCV crayfish compared with the random expectation (dotted line).

The median ED2 of a crayfish species is 10 MY, whilst the median ED2 of a CCV species is 12.3 MY. The highest median ED2 for a CCV species was 125 MY for Tenuibranchiurus glypticus, the second highest ranked EDGE2 species and smallest crayfish in the world at 3 cm, whilst the lowest was the Critically Endangered and visually striking Cherax tenuimanus with 0.433 MY.

For all four scenarios of projected CCV crayfish, the observed distributions of ED2 differed significantly (p < 0.0001) from the random expectations in Mann-Whitney tests (Fig. 5), with all scenarios except a 2070 intermediate warming affecting crayfish with higher ED2 scores more than expected and with each having large effect sizes. The greatest departure from the random expectation was for species projected to be CCV under a 2050 high warming scenario, where vulnerable species were on average 3.22 MY older (U = 962913, d = 0.926). This was followed by 2.2 MY older for a 2070 high warming scenario (U = 905521, d = 0.811) and then 1.77 MY for species projected as CCV under a 2050 intermediate warming scenario (U = 812544, d = 0.625). Species projected to be CCV under a 2070 intermediate warming scenario, however, had a lower ED2 than the random expectation by 0.76 MY (U = 331178, d = - 0.338).

We found that the simulated extinction of all 134 species listed as CCV across the four scenarios (median = 12,644 MY) resulted in 0.64% more PD being lost in comparison to the simulated extinction of the 86 crayfish that are listed as threatened by climate change using the Red List (median = 12,564 MY; U = 581982, p < 0.0001, d = 0.164). However, the crayfish threatened by climate change on the Red List had ED2 scores (median = 20.7 MY) that were 68.5% greater than those of the CCV crayfish (median = 12.3; U = 7834.5, p < 0.0001, d = 0.36).

## Discussion

Our study draws attention to the underappreciated plight of crayfish, a group where almost one-fifth of evolutionary history is at risk of extinction, and that are disproportionately threatened by climate change. Given that roughly one-third of crayfish lack sufficient conservation assessments, and that Data Deficient species are likely to have elevated extinction risks (Borgelt et al., 2022), this predicted loss is likely an underestimate. Furthermore, we highlight that threatened crayfish tend to be more evolutionarily distinct than non-threatened crayfish and, in our comparison with tetrapod animals, we demonstrated that crayfish rank amongst the most evolutionarily distinct clades examined so far.

Together, the 70 EDGE crayfish species identified here represent 40% of expected PD loss for all crayfish, despite comprising just a tenth of crayfish species richness. Conserving the remaining 70 threatened crayfish that do not qualify as EDGE species would safeguard just 10% more of the total expected loss. This demonstrates that safeguarding a relatively small number of priority species can protect a large portion of evolutionary history. If conservation efforts overlook evolutionary history as a key component of biodiversity, however, then we may face a situation where large and unique branches are lost from the Tree of Life (De Meester et al., 2024; Forest et al., 2015; Hendry et al., 2010).

Of the 70 EDGE priority species, 31 are within Australia’s largest crayfish genus Euastacus. This is the most threatened crayfish genus with more than one species, where 74% of the group’s 53 morphologically and ecologically diverse species are threatened with extinction (Furse & Coughran, 2011). The high proportion of EDGE species can be explained by the phylogenetic complementarity embedded within the EDGE2 metric, which ensures that closely related species that are similarly threatened have similar conservation value. In the case of Euastacus, the high concentration of priority species therefore reflects that each species represents a possible route to securing a portion of the whole clade’s shared evolutionary history. The EDGE priority list can then be used in accompaniment with known information on conservation feasibility to determine which species to protect and, in doing so, efficiently deliver huge gains for the preservation of the crayfish Tree of Life.

The nature of the EDGE metric explains the inclusion of and exclusion of other crayfish groups of interest. Tenuibranchiurus glypticus is a monotypic parastacid crayfish with the second highest ED2 score that is found in the swamps outside of Brisbane, a region of known value for its threatened tetrapod evolutionary history (Pipins et al., 2024). Given its extreme phylogenetic isolation and Endangered threat status, the species ranks as the second top scoring EDGE species on the list. Elsewhere, the Endangered Austropotamobius pallipes is Europe’s only EDGE crayfish. It appears on the EDGE list due to the genus’ notable evolutionary history, with the median genus-level ED2 score (29 MY) ranking sixth highest among all 34 genera, and the fact that it is the only member within its four species genus that has a data-sufficient Red List assessment. Some genera of interest do not feature on the list, like the genus Paranephrops, which contains the only two endemic crayfish on the islands of New Zealand, but which do not constitute conservation priorities as both species are marked as Least Concern on the Red List. Elsewhere, no species from the three South American endemic genera (Virilastacus, Samastacus, and Parastacus) make it on the list. This is because of a lack of conservation data on these genera, with all 4 Virilastacus species and the only Samastacus species being Data Deficient, along with 12 of the 14 Parastacus species, despite their high median ED2 scores (16 MY, 70.1 MY, and 19.7 MY, respectively) and the suggested threatened status of many (Almerão et al., 2015). Given its especially large ED2 score, Samastacus spinifrons was placed on the EDGE2 Research List, highlighting its urgent need for a conservation assessment. Elsewhere, no Malagasy species (Astacoides) make it on the list, despite four being Vulnerable (Crandall et al., 2022; Jones et al., 2007). This is due to their phylogenetic uncertainty with three of seven species requiring phylogenetic imputation. Three Astacoides species did make it on the EDGE Borderline list, reflecting their notable threatened evolutionary history which fails to meet the 95% certainty threshold.

Our study presents the first EDGE2 assessment for an invertebrate group, building on the previous phylogenetic crayfish prioritisation of Owen et al. (2015) but employing a standardised procedure that allows for comparisons to other EDGE2 assessed groups like tetrapod animals (Gumbs et al., 2024). Though the data deficiency observed in crayfish is beyond that of most tetrapod animal clades, we found that the metric still performed robustly, with species ranks remaining largely consistent before and after the imputation of the 35% of crayfish with absent phylogenetic data (Fig. 4). We therefore also highlight this study as being an important stepping stone for determining the validity of such methods for clades with even larger data gaps, such as in flowering plants, fungi, and insects.

Our results echo the concerns of previous studies on the impact that climate change will have on phylogenetic diversity (Faith & Richards, 2012; Gonzalez-Orozco et al., 2016). A high warming scenario could affect up to 409 million more years of crayfish evolutionary history than expected at random, whilst following an intermediate warming scenario could spare 251 MY in comparison. Furthermore, a high warming scenario threatens individual species that are more evolutionarily distinct than the random expectation by 3.22 MY, due to the greater proportion of parastacid crayfish that are considered CCV in high warming scenarios (Hossain et al., 2018). The parastacids, native to southern latitudes, are known to have lower diversification rates than crayfish of northern latitudes and exhibit longer terminal branch lengths (Owen et al., 2015; Toon et al., 2010). Australia is the centre of parastacid diversity and, owing to their extensive evolutionary history, parastacids are also disproportionately represented as EDGE species, making up 44 of the 70 EDGE priority crayfish, with 23 of these being climate change vulnerable (Hossain et al., 2018). Avoiding a high warming scenario will therefore be essential for the conservation of Australia’s highly distinct crayfish.

We found that only 29 species were jointly marked as being at risk from climate change according to the climate change trait-based vulnerability analysis (CCVA) and the Red List threat classification. This is unsurprising since IUCN Red List assessments generally focus on a shorter time period than that evoked by CCVA. There may also be a bias inherent in Red List assessments, which often generalize associations, for example, between temperature tolerance and climate change risk to generate assessments for large species groups (Trull et al., 2018). Meanwhile, trait-data incompleteness introduces ambiguity; in the crayfish CCVA of Hossain et al. (2018), traits with unknown trait values were scored as low in relation to a species’ overall vulnerability to climate change, leading to an average of 18% of recorded crayfish being listed as CCV. In contrast, when unknown trait values were instead scored as high, an average of 48% of species were CCV. Approximately one-third of the crayfish used in this study had no information relating to their risk from climate change. When these discrepancies are taken into consideration, it is possible that a much larger proportion of crayfish PD is threatened than our study suggests.

Phylogenetic diversity has been gaining traction within conservation, with recent developments including the EDGE index featuring as a complementarity indicator under Target 4 of the Kunming-Montreal Global Biodiversity Framework (Secretariat of the Convention on Biological Diversity, 2022) and the establishment of the IUCN SSC Phylogenetic Diversity Task Force (Gumbs et al., 2023a). We believe the creation of this crayfish EDGE list can capitalise on this momentum by feeding into the conservation activities of the EDGE of Existence programme, which to date has funded work on over 130 EDGE species. Furthermore, although an increasing number of crayfish are being recognised by the Endangered Species Act in the US (Taylor et al., 2019), crayfish are still rarely featured in freshwater conservation policy and action globally (Danilović et al., 2022; Taylor et al., 2019). As most EDGE crayfish and climate change vulnerable crayfish are located within Australia, we advise a reassessment of this country’s threatened parastacids coupled with action to secure their long-term future, taking phylogenetic diversity into account across the bioregionalization of Australia (Whiting et al., 2000).

Given that a high warming scenario threatens a disproportionately large amount of crayfish PD and evolutionarily distinct species, the absence of climate mitigation strategies will likely see the loss of some of the world’s most unique and irreplaceable crayfish. If action can be taken to curb carbon emissions and avoid a worst-case scenario, however, we could safeguard hundreds of millions of years of crayfish evolutionary history.

## Supporting Information

Additional supporting information may be found online in the Supporting Information section at the end of this article.

## Acknowledgements

SP was funded by the NERC Science and Solutions for a Changing Planet Doctoral Training Programme (grant number NE/S007415/1), the CASE component of which was funded by On the Edge. MB was supported by a grant from the Rufford Foundation. KAC was supported as a VisitANTS Visiting Scholar as part of the Biodiverse Anthropocenes program at the University of Oulu, Finland.

## Author contributions

RG, JR, and MB conceived the study. SP, RG, and JR designed the analyses. SP conducted the analyses. LB and MAH provided crayfish trait-based vulnerability analysis data. KAC provided crayfish phylogenetic data. RG provided the EDGE2 code and tetrapod data. RG, JR, and MB supervised the study. SP led the writing of the manuscript. All authors contributed critically to the drafts and gave final approval for publication.

