## Appendix S1 for "Phylogenetically-informed crayfish conservation in the face of climate change"

**Supplementary Information**

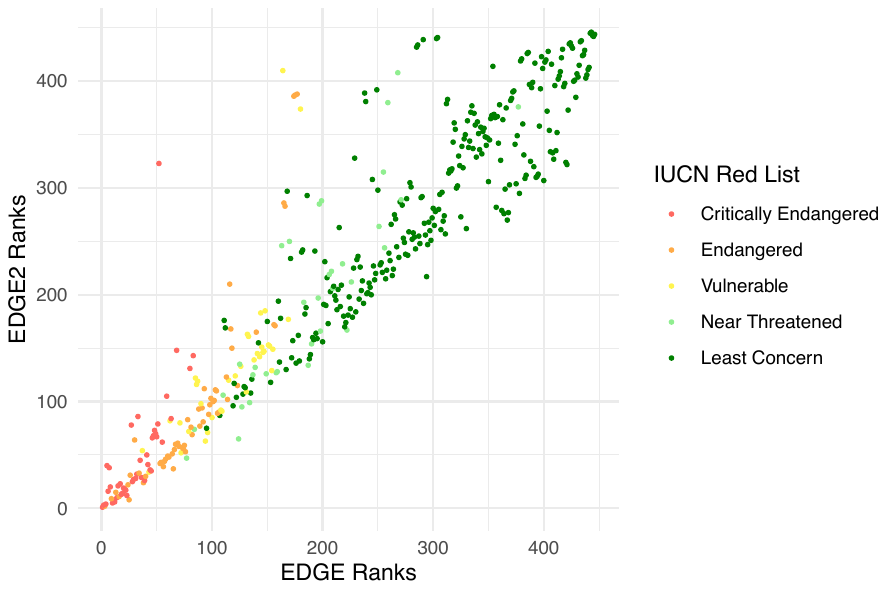

**Appendix S1:** A comparison of species ranks from EDGE and EDGE2, where points represent individual crayfish species and are coloured by their IUCN Red List categories.
